# Loss of α2-6 sialylation promotes the transformation of synovial fibroblasts into a pro-inflammatory phenotype in Rheumatoid Arthritis

**DOI:** 10.1101/2020.03.08.970046

**Authors:** Yilin Wang, Aneesah Khan, Aristotelis Antonopoulos, Laura Bouché, Christopher D Buckley, Andrew Filer, Karim Raza, Kun Ping-Li, Barbara Tolusso, Elisa Gremese, Mariola Kurowska-Stolarska, Stefano Alivernini, Anne Dell, Stuart M. Haslam, Miguel A. Pineda

**Affiliations:** Institute of Infection, Immunity and Inflammation, University of Glasgow, UK; Department of Life Sciences, Imperial College London, London, United Kingdom; Rheumatology Research Group, Institute for Inflammation and Ageing, College of Medical and Dental Sciences, University of Birmingham, Queen Elizabeth Hospital, Birmingham, United Kingdom; The Kennedy Institute of Rheumatology, University of Oxford, Oxford, United Kingdom; Sandwell and West Birmingham Hospitals NHS Trust, Birmingham, United Kingdom; School of Pharmacy, Guangdong Pharmaceutical University, Guangzhou, China; Division of Rheumatology, Fondazione Policlinico Universitario A. Gemelli IRCCS, Rome, Italy; Division of Rheumatology, Università Cattolica del Sacro Cuore, Rome, Italy; Research into Inflammatory Arthritis Centre Versus Arthritis (RACE), Glasgow, Birmingham, Newcastle, Oxford, United Kingdom

## Abstract

In healthy joints, synovial fibroblasts (SFs) provide the microenvironment required to mediate homeostasis but are recognized to adopt a pathological role in rheumatoid arthritis (RA), promoting the infiltration and activation of immune cells to perpetuate local inflammation, pain and joint destruction. Carbohydrates (glycans) attached to cell surface proteins are fundamental regulators of cellular interactions between stromal and immune cells, but very little is known about the glycome of SFs or how glycosylation regulates their biology. Here we fill these gaps in our understanding of stromal guided pathophysiology by systematically mapping glycosylation pathways in healthy and arthritic SFs. We used a combination of transcriptomic and glycomic analysis to show that transformation of fibroblasts into pro-inflammatory cells in RA is associated with profound glycan remodeling, a process that involves reduction of α2-6 terminal sialylation that is mostly mediated by TNFα-dependent inhibition of the glycosyltransferase ST6Gal1. We also show that sialylation of SFs correlates with distinct disease stages and SFs functional subsets in both human RA and models of mouse arthritis. We propose that pro-inflammatory cytokines in the joint remodel the SF-glycome, transforming a regulatory tissue intended to preserve local homeostasis, into an under-sialylated and highly pro-inflammatory microenvironment that contributes to an amplificatory inflammatory network that perpetuates chronic inflammation. These results highlight the importance of cell glycosylation in stromal immunology.

## Introduction

Rheumatoid Arthritis (RA) is a chronic inflammatory condition of the joints affecting 0.3%-1% of the world’s population. RA has been historically described as an autoimmune condition, where the central component of disease pathogenesis relies on aberrant responses of immune cells leading to destruction of bone and cartilage in the joint. Recent findings have now shown that the pathophysiology of RA merges autoreactive immunity with genetic, epigenetic and environmental factors that are responsible for disease initiation. RA starts with a pre-clinical phase involving activation of immune mechanisms in the absence of clinical symptoms^1^, followed by auto-amplificatory loops attracting macrophages, T cells and other immune cells to the joint to perpetuate local inflammation through cytokine networks dominated by TNFα, IL-6, IL-1β and chemokines such as Ccl2 or CXCL5^2,3^. Not surprisingly, biologic Disease Modifying Anti-Rheumatic Drugs (bDMARS) inhibiting these cytokines are the treatments of choice in the clinic. Even though such gold standard treatments lead to a substantial improvement in life quality of thousands of patients^1,4^, as they are immunosuppressive agents they have been linked with serious adverse effects. Also 30-40% of RA patients still do not respond completely to them, suggesting that key events underpinning pathogenesis remain elusive. In fact, the cellular and molecular basis of why inflammation does not resolve in RA remains unanswered.

Synovial fibroblasts (SFs). are major components of the synovial membrane, a highly specialized mesenchymal tissue lining the joint cavity. In the synovium, two main micro domains can be described; the lining layer (directly exposed to the synovial space) and sublining layers. Due to their anatomical location, SFs provide the required nutritional and structural support. Initially SFs were considered as cells lacking any substantial impact on immune function. However SFs are known to adopt a immunopathological role in RA, responding to inflammatory cytokines and promoting bone and cartilage destruction^2,3,5,6^. Despite being cells of non-immune origin, SFs play a central role to perpetuate local immune responses in the synovial joint, delivering region-specific signals to infiltrating immune cells and contributing to bone and cartilage destruction^7,8^. Interestingly, it has been recently demonstrated by Single Cell Transcriptomics that SFs comprise distinct functional subsets that correlate with their anatomical location and activation of pathological pathways^9,10^, suggesting that SF-dependent immunity may be far more complex than anticipated. Because of their non-immune origin and their highly-specialized function in health and disease, interventions targeting SFs -or specific SFs subsets-may modify disease progression without significant immunosuppression.

Functional glycomics, an emerging discipline focused on defining the structures and functional roles of carbohydrates (glycans) in biological systems, could offer such fibroblast specific molecular targets^11,12^. Some elegant studies have shown the potential impact of glycan regulation in multiple aspects of RA pathophysiology, for example reduced sialylation of N-glycans is a feature of pathogenic immunoglobulins in RA patients and their glycosylation profile shows predictive potential for disease progression ^13-17^. Besides, galectins, a family of proteins that bind to galactose-containing glycans, are key modulators of synovial inflammation^18-20^. However, little is known about the glycosylation profile of SFs, and whether this varies in health and disease, despite the glycome is being critical for dictating interactions with galectins and other carbohydrate binding proteins, and the fact that altered glycosylation is a hallmark of chronic inflammatory conditions. To illustrate this, cytokines induce aberrant glycome changes in cancer that perpetuate local inflammation via control of cell adhesion, migration and signal transduction^21-23^; mechanisms that are also associated to the pathogenicity of SFs in RA and are responsible of migration to cartilage-containing areas prior to tissue damage. Furthermore, the carbohydrate binding protein galectin-3 is up-regulated in RA^24^ and galectin-3^-/-^ mice show reduced pathology in experimental arthritis^20^. Moreover, exogenous galectin-3 significantly up-regulates CCL2, CCL3 and CCL5 in synovial but not in dermal fibroblasts^19^, suggesting that the synovium microenvironment can induce tissue-specific glycosylation in the stromal compartment.

We hypothesised that the cytokine milieu found in the inflamed joint determines distinct SF-glycosylation, that in turn, regulates cell recruitment and inflammatory responses. Our aim in this study was to investigate changes of SF glycome that could be related to their inflammatory activity. By combining transcriptomics and glycomics analysis, we report that transformation of SFs into pro-inflammatory cells in RA is associated with glycan remodeling, involving reduction of terminal sialylation in Thy1(CD90)^+^ sublining SFs upon TNFα stimulation. We also show that low sialylation of SFs is associated with disease remission, supporting the idea that the stromal glycome could be used for development of novel disease biomarkers or therapeutic agents.

## Results

### Distinct anatomical locations and inflammatory conditions shape the glycosylation profile of human fibroblasts

To test our initial hypothesis, we isolated and expanded fibroblasts from the synovium and matched samples from skin from RA patients, as a reflection of inflammatory and non-inflammatory environments. We also isolated SFs from osteoarthritis (OA) patients, as an example of a less inflammatory, more destructive joint disease. General glycosylation was evaluated using lectin-binding assays by immunofluorescence (Fig. 1a) and flow cytometry (Fig. 1b). Based on the carbohydrate binding specificity for each lectin, our results provided an initial indication of differential fibroblast glycosylation under inflammatory conditions. All fibroblasts bound most of the tested lectins, confirming the presence of a rich and diverse glycocalix. These data indicate the SF-glycome was rich in galactose-containing glycans (RCA^+^, ECA^+^) containing PolyLacNAc extensions (LEL^+^) and α1-6 core fucosylated N-glycans (AAL^+^), in contrast to the lack of α1,2 fucosylation on glycan antennae (UEA^-^). Interestingly, we observed significant differences in glycosylation in fibroblasts taken from distinct anatomical locations (dermal vs synovium: PNA, Jacaline and ECA binding) and in distinct inflammatory conditions (RA vs OA, LEL and ECA binding) (Fig. 1a-b), supporting the link between local inflammatory mediators and glycan remodeling. To evaluate whether changes in glycosylation were due to cytokine stimulation, we checked the expression of glycan building enzymes, glycosyltransferases, in human SFs in response to pro-inflammatory TNFα [dataset generated by Slowikowski et al.^25^]. TNFα significantly modulated glycosyltransferase genes involved in the synthesis of branched glycans such as A4GALT, GCNT1 and GCNT2, along with up-regulation of α2-3 sialyltransferases like ST3Gal1, ST3Gal2 and ST3Gal4 (Supp. Fig. 1a). No effect was observed when cells were stimulated with IL-17 (Suppl. Fig. 1b), indicating that changes in glycosylation are linked to individual cytokine signaling. To further explore these findings, naive SFs expanded from mouse synovium were stimulated with a panel of regulatory and pro-inflammatory factors and subsequently stained with PHA, lectin that binds complex branched N-glycans.

**Figure 1:**
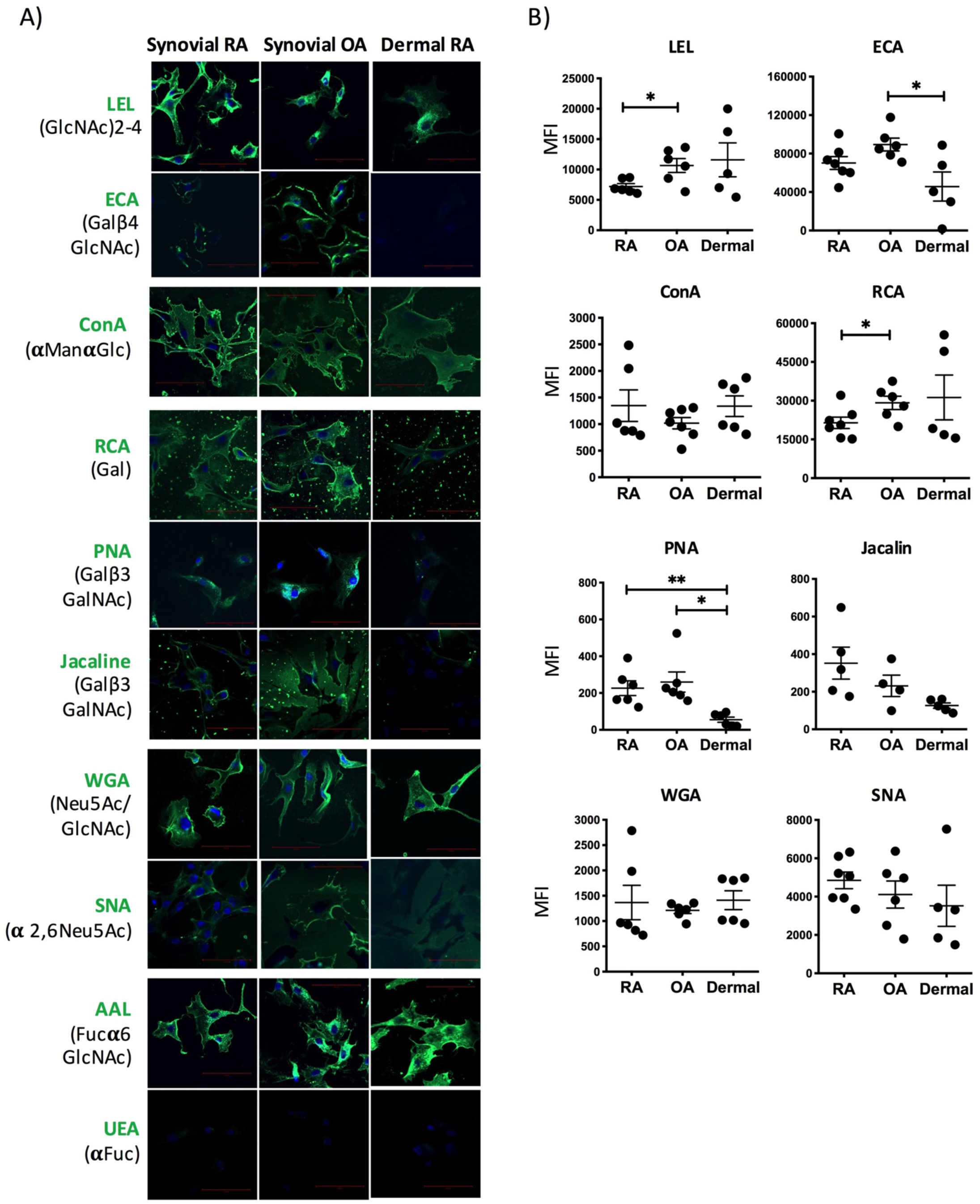
Lectin staining shows that glycosylation of human fibroblasts varies in distinct anatomical locations or in inflammatory environments. **a)** Immunofluorescence staining (lectin, green) and nuclei (DAPI, blue) of synovial and dermal fibroblasts isolated from RA patients following joint replacement surgery, and synovial fibroblasts isolated from OA patients. Lectins and their carbohydrate binding specify are shown. Scale bars = 100 μm. **b)** Flow-cytometric analysis of indicated lectin staining on non-permeabilised synovial fibroblasts (RA) and dermal fibroblasts (Dermal) from RA patients, and synovial fibroblasts from osteoarthritis patients (OA). Data show the Mean Fluorescence Intensity (MFI) of the whole population. Statistics: all results are presented as mean ± SEM and each symbol represents one individual patient (RA, n=6-7, OA, n=4-7, Dermal, n=5-6). Statistical significance was determined using Brown-Forsythe and Welch ANOVA test for multiple comparisons and significance indicated by asterisks, * p < 0.05 and ** p < 0.01.

The cytokine IL-22 was chosen because it exerts both inflammatory and resolving actions in RA, depending on the local microenvironment^26^. Differential effects of IL-22 over cell glycome would therefore provide further support to the original hypothesis linking distinct glycosylation and effector responses. The combination of inflammatory factors with IL-22 exerted diverse effects on expression of branched glycans (Suppl. Fig. 1c), perhaps related to the dual effect described by IL-22 in joint inflammation. Changes in PHA-binding in response to immunomodulators might be reflective of SFs activation as glycans recognized by PHA were detected in joint areas with a strong SF-mediated inflammation (Suppl. Fig. 1d). Overall, results shown in Fig. 1 and Suppl. Fig. 1 supported the hypothesis that immune mediators found in local microenvironments control SF glycosylation.

### Comparative RNA-Seq analysis reveals that transformation of SFs into pathogenic mediators is associated with changes glycosylation pathways

To complement the lectin-based experiments we additionally undertook transcriptome sequencing (RNA-Seq) to fully dissect SFs glycosylation pathways at mRNA level. We also chose to work initially with the murine model of Collagen-Induced Arthritis (CIA), because human clinical samples have usually been exposed to long-term immunomodulatory treatments, which add high levels of variability and might affect the cytokine-glycosylation axis. The CIA model shares hallmarks of human disease of high relevance to this study, such as the hyperplasia of synovial membrane, inflammatory infiltration of the synovium and pannus formation. SFs from healthy and CIA mice (Clinical scores >8) were sorted by flow cytometry [CD45^-^ CD31^-^Podoplanin^+^] (Fig. 2a) and RNA was immediately purified for RNA-Seq analysis upon polyA selection. We recovered a higher number of SFs from CIA joints compared to healthy joints with elevated expression of podoplanin (Fig. 2a), reflecting their hyperplasia and activation reminiscent of human RA. Consistently, Principal Component Analysis of transcriptomic data confirmed the distinction between both groups (Fig. 2b). A list of differentially expressed (DE) genes in CIA SFs was generated including 298 up-regulated genes and 88 down-regulated [>2 fold, adjp <0.05, (Fig. 2b, suppl. Fig. 2a)]. KEGG pathway enrichment analysis showed that ‘Rheumatoid Arthritis’ as disease pathway was significantly up-regulated in CIA SFs (Suppl. Fig. 2b), validating the model to study SF-dependent inflammation. String Protein-Protein Interaction Networks Functional Enrichment Analysis^27^ was applied to DE genes in CIA compared to healthy SFs, identifying 2 main functional networks: i) cell cycle and cell division and ii) inflammatory response (Fig. 2c), further demonstrating SF immune activation and hyperproliferation. Interestingly, GO-term analysis revealed that most proteins identified in the inflammatory network were glycoproteins and/or regulators of cell communication (Fig. 2c), suggesting the potential role of glycosylation in SF activation. To further study this, we specifically searched for expression of genes involved at different steps of glycosylation biosynthesis, such as glycosyltransferases and glycosidases responsible of mannosylation, glycan chain branching and elongation, fucosylation, sialylation and glycan degradation (Fig. 2d). Unsupervised clustering based on these glycan biosynthetic pathways separated naïve and arthritic SFs (Fig. 2e), suggesting that pro-inflammatory SFs present an altered glycome in inflammatory conditions. The observed down-regulation of enzymes of medial Golgi-branching N-acetylglucosaminyltransferases II, IV and V (encoded by Mgat2, Mgat4 and Mgat5) in CIA SFs, along with the up-regulation of β-1,4-Galactosyltransferase genes (B4GALT) and GCNT2 (Fig. 2e) suggested that these differences might lead to changes in the extension and branching of antennae of N-glycans or in the number of poly N-acetyllactosamine (linear repeats of Galβ1,4GlcNAcβ1,3) by β1,3N-acetylglucosaminyl-transferases. Likewise, terminal modifications of such structures may have reduced fucosylation or sialylation, given that gene expression of the fucosyltransferases Fut10, Fut11 and the sialyltransferases ST6Gal1, ST3Gal2 and ST3Gal6 are significantly down-regulated in CIA SFs (Fig. 2e).

**Figure 2:**
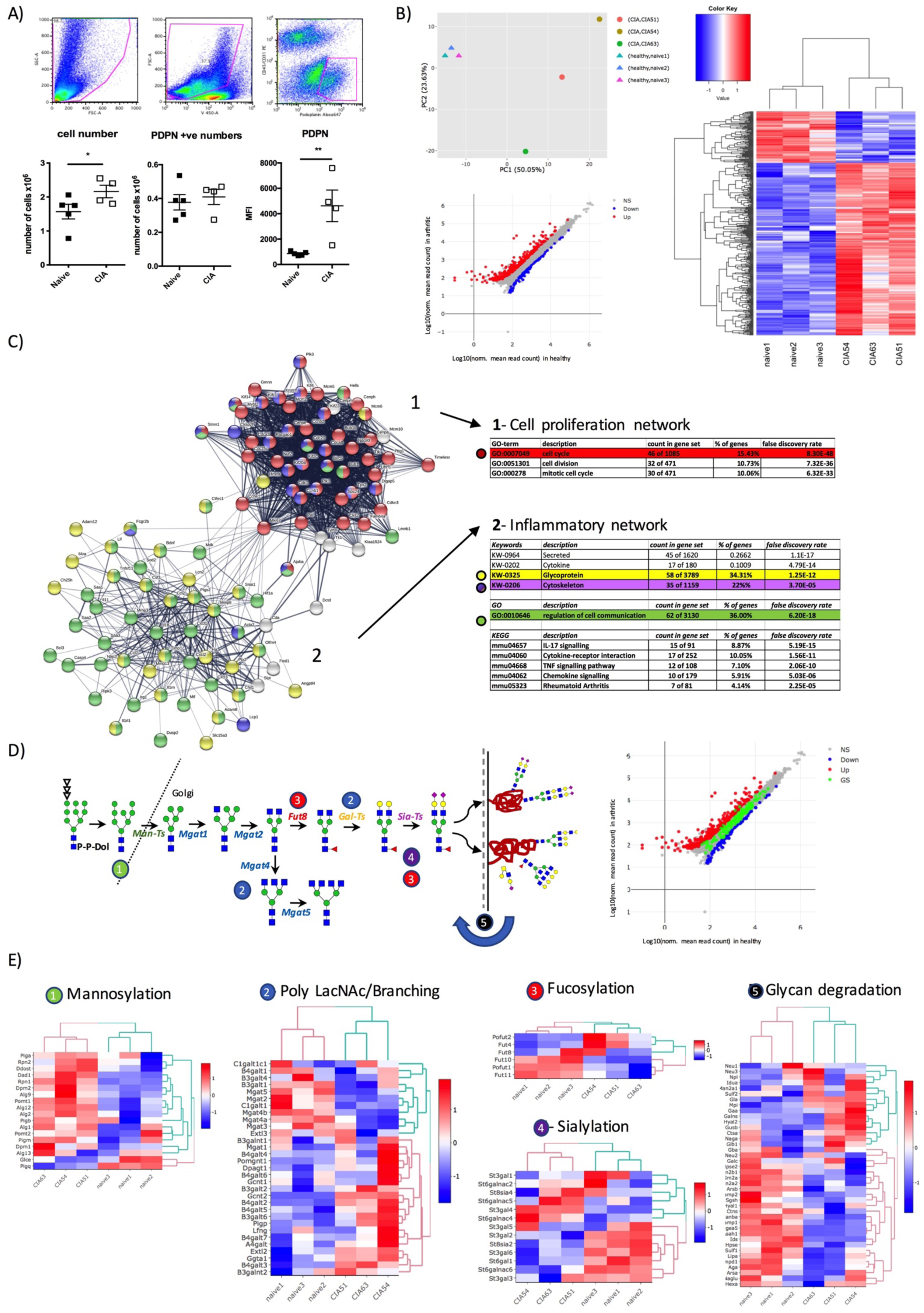
RNA-Seq studies confirm the unique transcriptomic profile of glycosylation pathways in inflammatory synovial fibroblasts. Paws from healthy and arthritic mice were dissected and soft tissue was digested. **a)** Live synovial fibroblasts (Zombie Violet^-^, Podoplanin^+^, CD45^-^, CD31^-^) were sorted by flow cytometry as shown in dot plots. Number of isolated cells, PDPN+ and expression of PDPN is show. Naïve n=5, arthritic n=4. Statistical significance was calculated using a one-tail unpaired t test, *p < 0.05 and **p < 0.01. **b)** RNA was isolated (RIN>9) from healthy (n=3) and arthritic CIA (n=3, scores of 9,10 and 11) mice and subjected to bulk RNA-Seq (75bp paired-end, 30M reads). Principal component analysis (PCA) and differential expression (DE) of genes are shown. All detected genes are plotted as a scatter plot where x=gene expression healthy, y=gene expression arthritic. Genes that pass a threshold of padj < 0.01 and |log2foldChange| > 2 in DE analysis are colored in blue when they are down regulated and red when they are upregulated in the arthritic (CIA) mice. Heatmap shows up and down-regulated genes; [unsupervised clustering in rows and columns based on Euclidean distances] **c)** Function enrichment and network analysis regulated by synovial inflammation. STRING protein-protein interaction network (https://string-db.org) was performed on DE genes from (b). Significantly modulated pathways and cellular components associated to DE genes in arthritic mice are shown in tables. [PPI enrichment p-value: < 1.0e-16]. K-Means method gave two main functionally related clusters of genes, designated as ‘cell proliferation’ and ‘inflammatory’ cluster. Color code for nodes are: red: *cell cycle*, purple: *cytoskeleton*, yellow: *glycoprotein*, green: *regulation of cell communication*. **d)** Analysis of glycogenes and glycan biosynthesis pathways. Illustration shows N-glycan synthesis pathway. Scatter blot: as in (b) but with genes annotated in the glycan biosynthesis pathways highlighted in green. **e)** Heatmaps of scaled and centered Log_10_ transformed normalized read counts showing only differentially expressed genes involved in glycosylation pathways.

### Glycomic studies reveal reduced levels of sialylation in pro-inflammatory SFs

Transcriptomic analysis provided a powerful tool to delineate potential changes in cell glycosylation. However, unlike proteins or nucleic acids complex, glycans are not assembled in a template-driven process. Rather, glycosylation in the endoplasmic reticulum and Golgi is the result of combined actions of glycosyltransferases and glycosidases (Fig. 2d,e). Consequently, prediction of structures based on transcriptomic data does not necessarily correlate with the final glycosylation profile and further structural information is needed to generate reliable information. We therefore used mass spectrometry (MS) based glycomics to define the N-glycome of murine SFs (Suppl. Fig. 3). N-glycans were isolated from cultured SFs, permethylated and subjected to MS analysis. Following annotation of peaks with most likely glycan structures based on molecular ion composition, knowledge of biosynthetic pathways and with the assistance of the bioinformatic tool glycoworkbench ^28^, most structures were annotated as high mannose glycans or complex glycans, either core-fucosylated or non-fucosylated. Sequential addition of N-acetyllactosamines (LacNAcs) defined the larger structures, suggesting the presence of extended antennae and multi-branched structures. Sialylation was the most abundant capping modification of terminal galactoses. As expected in murine cells, two sialic acids were detected, N-acetylneuraminic and N-glycolylneuraminic acid. A similar structural profile was observed in human SFs taken from RA joint replacements (Suppl. Fig. 4), further validating the animal model. Still, some differences between mouse and human cells were found, like the absence of terminal Gal-α-gal and N-glycolylneuraminic acid, as these structures cannot be synthesized by the human glycosylation machinery. Nevertheless, human and mouse SF N-glycomes seemed to be well conserved.

**Figure 3:**
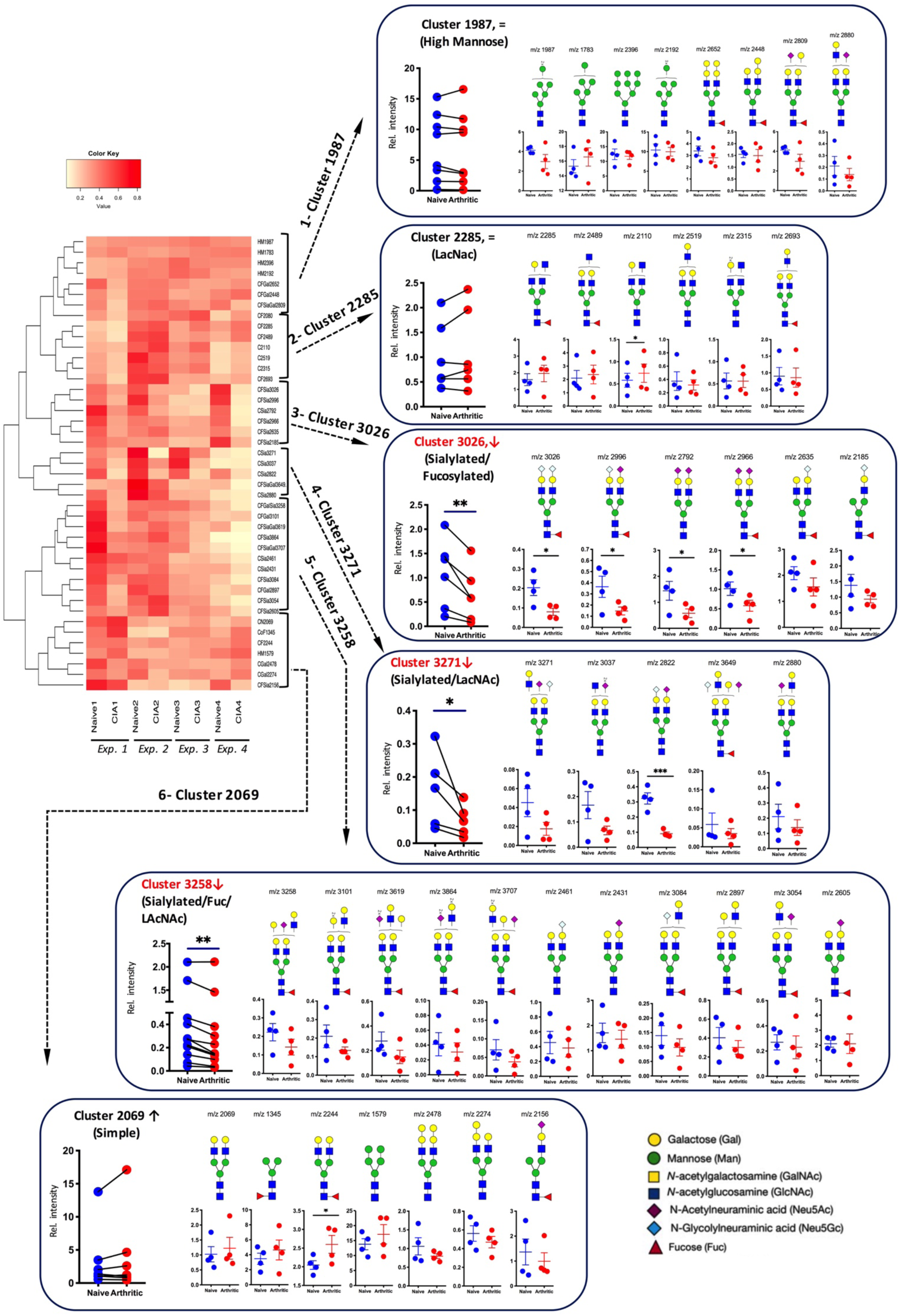
Glycomic studies in experimental Collagen-Induced Arthritis show a reduction in sialylated N-glycans in synovial fibroblasts associated to inflamed joints. N-glycans from synovial fibroblasts expanded from healthy or arthritic synovial fibroblasts were isolated and permethylated prior to MALDI-TOF MS analysis. 43 structures were selected (relative expression >2%) and MS peak area were quantified and normalized against total measured intensities. Unsupervised hierarchical clustering with Euclidean distance grouped structures into six main clusters as shown. Structures present in each cluster are shown. In all graphs, blue dots represent healthy synovial fibroblasts and red dots are cells from CIA animals. Data show the results from 4 independent experiments, where healthy and arthritic fibroblasts samples were processed in parallel. Before-after graphs show all structures present in the indicated clusters, where individual dots are the mean of relative expression from 4 experiments and similar structures in healthy and CIA cells are connected with lines. Statistical significance between healthy and CIA was assessed by one tail paired t-test, *p < 0.05 and **p < 0.01. Relative expression for each glycan structure (shown with their *m/z* value) is also shown in scatter plots showing mean and SD, where each dot represents the relative expression of one independent experiment, n=4, *p < 0.05 and **p < 0.01 using unpaired one tail t-test.

**Figure 4:**
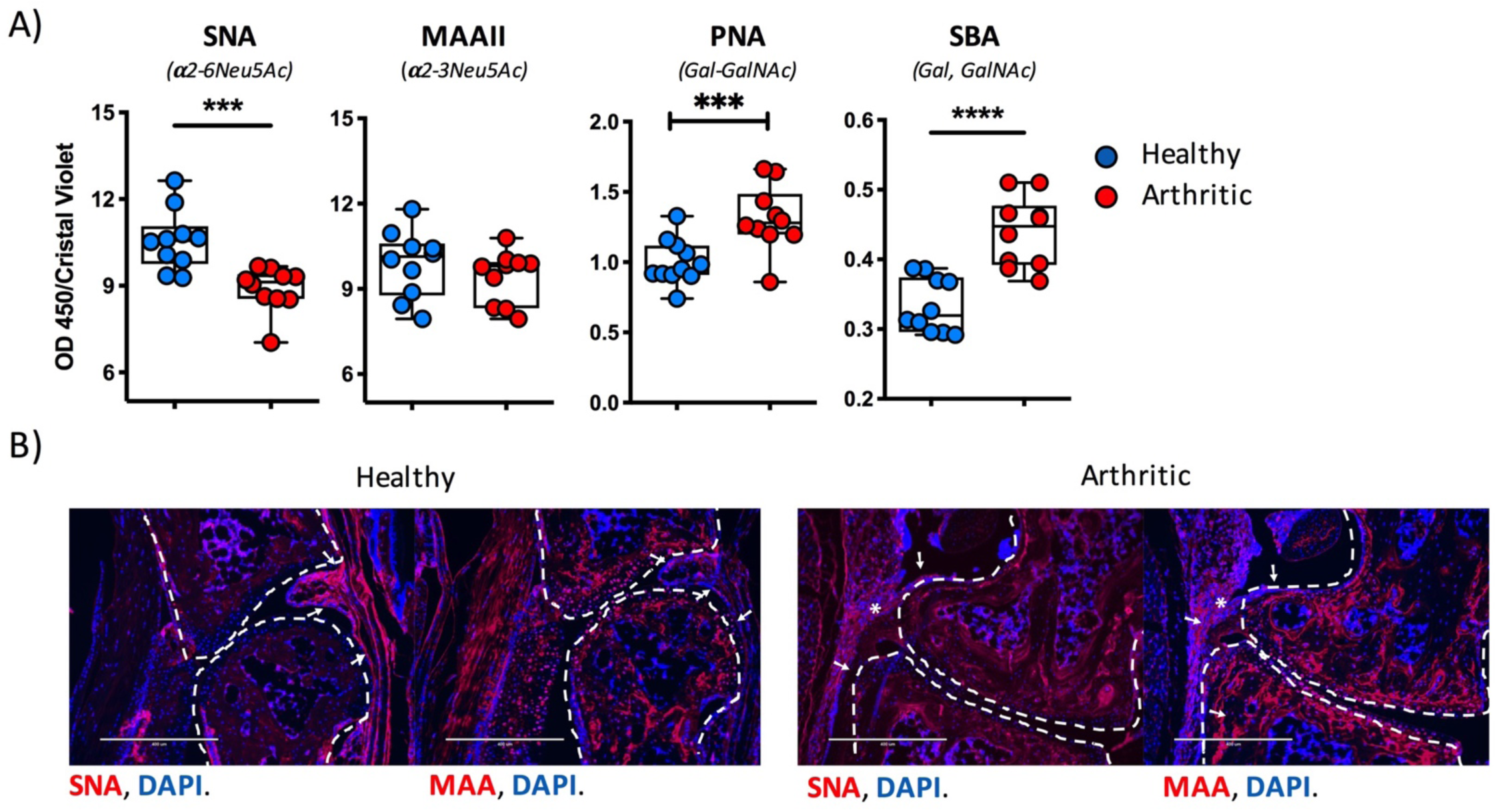
Synovial fibroblasts from arthritic mice show reduced levels of *α*2-6 but not *α*2-3 sialylation. **a)** SNA, MAAII, PNA and SBA lectins were used in Enzyme-linked lectin assay (ELLA) to evaluate the presence of *α*2-6 Sialic acid, *α*2-3 sialic acid, non-sialylated T antigen and terminal N-acetylgalactosamine/galactose respectively. Statistics: unpaired t-test, n=10. *p < 0.05, **p < 0.01, ***p < 0.001. b) Mouse joint sections were stained with biotinylated SNA and MAAII and fluorescently labelled streptavidin. Magnification 20x, scale bars: 200 μm. Lectin binding is shown in red, nuclei counterstain with DAPI in blue. Superimposed white elements indicate as follows; [dotted lines]: bone limits, [arrows]: synovial membrane, [asterisks]: local inflammation and infiltrating cells.

Next, in order to identify specific glycan changes that could contribute to SF activation, the relative expression of individual N-glycans structures in healthy and CIA SFs were compared. Structures whose relative expression was below 2% of total were ruled out, since the low expressed forms could easily add artefactual results, leaving 43 glycans to analyze. This dataset was then used to perform unsupervised hierarchical clustering to uncover expression patterns characteristic of inflammatory CIA SFs (Fig. 3). Interestingly, N-glycans were clustered into 6 discrete groups containing similar structural features: i) *cluster 1987*: high mannose, ii) *cluster 2285*: LacNAc extended, iii) *cluster 3026*: sialylated-fucosylated, iv) *cluster 3271*: sialylated-LacNAc v) *cluster 3258*: sialylated-LacNAc-fucosylated, and vi) *cluster 2069*: simple non-sialylated. Three of these clusters contained glycans that were significantly down-regulated in CIA SFs: clusters *3026, 3271* and *3258*, which had one characteristic in common: a significant high proportion of sialic acid containing glycans. In fact, 96% of all the sialylated glycans analyzed were found within these three clusters, strongly suggesting that reduced sialylation is associated with activated CIA SFs.

### SFs from arthritic mice exhibit down-regulated α2-6-linked sialic acid

Comparative glycomic analysis (Fig. 3) indicated that experimental arthritis was strongly associated with a down-regulation in sialic-acid-containing N-glycans in SFs. Retrospective examination of the data related to transcriptomic regulation reveals a significant down-regulation of ST3Gal6, ST3Gal2 and ST6Gal1 [fold-increase 0.73, 0.64 and 0.60 respectively] (Fig. 2e), supporting the reduced presence of sialylated glycans observed by MS (Fig. 3). However, it also showed a significant up-regulation of ST3Gal4 mRNA [1.49 fold-increase (Fig. 2e)]. These apparently conflicting results could be explained by differential regulation of sialic acid linkages, differences that would not be detected in our initial MS-based studies. In the structures depicted in Fig.ure 3, two major types of glycosidic bonds can be formed between sialic acid and terminal galactose, namely, sialic acid-α2-3Gal and sialic acid-α2-6Gal, synthesized by six ST3 beta-galactoside alpha-2,3-sialyltransferases (ST3Gal1-6) and two ST6 beta-galactoside alpha-2,6-sialyltransferases (ST6Gal1-2). To evaluate the type of sialylation present in arthritic SFs, we used SNA and MAAII, lectins that specifically recognize sialic acid in α2-6 and α2-3-linkages. Results showed that SNA binding was reduced in CIA SFs compared to naïve SFs, whereas MAAII binding was not affected (Fig. 4a). These results indicate that the differential sialylation profile observed in CIA SFs is due to specific reduction in α2-6 sialylation, presumably because of lower ST6-sialyltransferases expression. Furthermore, binding of galactose-recognizing lectins such as PNA and SBA was up-regulated in arthritic SFs (Fig. 4a), probably reflecting increased terminal galactose in the under-sialylated glycome. Since inflammation does not change α2-3-sialylation of SFs, a reduced ratio of α2,6/α2,3-linked sialic acid might constitute a hallmark of inflammatory SFs. Supporting these findings, synovial membranes of healthy mouse joints had a α2-6 > α2-3 sialylation profile, as observed by immunofluorescence with SNA and MAAII, whilst inflamed joints in CIA joints show comparable levels of both sialic acid linkages (Fig. 4b).

### Down-regulation of α2-6 sialylation in SFs is a consequence of TNFα-dependent inhibition of the sialyltransferase ST6Gal1

The results from transcriptomics, MS-based glycomics and lectin-binding experiments all concluded that reduced α2-6**-**sialylation is associated with inflammatory SFs. We therefore next sought to identify the molecular mechanisms underlying this shift in the SF-glycome. Several sialytransferases (Sia-Ts) are expressed in SFs (Fig. 5a), including ST3Gal enzymes [ST3Gal1>>ST3Gal2>ST3Gal4>ST3Gal3] and only one involved in α2-6-Sialylation, ST6Gal1, with some significant gene regulation during CIA as explained before. ST8Sia-T involved in polysialic synthesis were only marginally expressed, in agreement with MS glycomics data (Suppl. Fig. 3). In addition to Sia-Ts, we studied expression of enzymes synthesizing their substrate, CMP-Neu5Ac (Fig. 5a), since a reduction in its intracellular concentration would also lead to hyposialylated glycomes, even if the expression of Sia-Ts remained at normal levels. As expected, given the abundant sialic acid content of SFs glycoproteins, these enzymes were highly expressed in both healthy and CIA SFs (Fig. 5a), suggesting that intracellular availability of Sialic acid donors might not restrain glycan biosynthesis. However, and rather unexpectedly, neither RNA-Seq analysis nor qPCR approaches detected expression of N-Acetylneuraminic Acid Phosphatase (NANP), an enzyme that dephosphorylates sialic acid 9-phosphate to free sialic acid as part of the required intermediate biosynthetic metabolites. As sialylated glycans are undeniably present in SFs, alternative biosynthesis pathways might be operating in these cells, perhaps as in recent observations in CHO cells^29^. To identify possible mechanisms behind changes in sialylation, we used qPCR to evaluate expression of selected genes (as in Fig. 5a) in response to IL-1β, IL-17 and TNFα, key cytokines known to initiate inflammatory pathways in RA. IL-6 mRNA, included as a positive control, was significantly up-regulated in response to all cytokines tested (Fig. 5b), confirming cell activation. No difference was observed in GNE, NANS, CMAS or SLC35A1 expression, corroborating our findings that sialic acid biosynthesis is not the cause for the reduced sialylation observed in CIA SFs. Regarding SiaTs expression, no effect was seen in response to IL-17, in line with the human observations in Suppl. Fig. 2b. IL-1β increased expression of ST6Gal1 (in Naïve SFs) and ST3Gal4 (in CIA SFs), with an approximately 2-fold increase. However, these changes were only mild compared with the significant down-regulation of ST6Gal1 in response to TNFα, that induced a strong down-regulation of ST6Gal1 (8-fold) in both naïve and CIA SFs (Fig. 5b). Since ST6Gal2 was not detected in SFs, TNFα-mediated down-regulation of ST6Gal1 seems to a key mechanism responsible for reduced SF sialylation in CIA arthritic mice. To confirm the functional effect over the SF-glycome, we repeated the experiment evaluating in parallel the expression of α2-6-sialic acid (SNA^+^ binding). Again, only TNFα decreased ST6Gal1 mRNA and such effect was accompanied by reduction of α2-6-Sialic acid on the cell surface (Fig. 5c). Finally, cells were stimulated with TNFα for 48 hours to allow glycan turnover, and N-glycans were then isolated to conduct MS-based glycomic analysis. 12 out of 16 sialylated N-glycans showed a reduced expression upon TNFα stimulation, providing further support to our findings (Fig. 5d).

**Figure 5:**
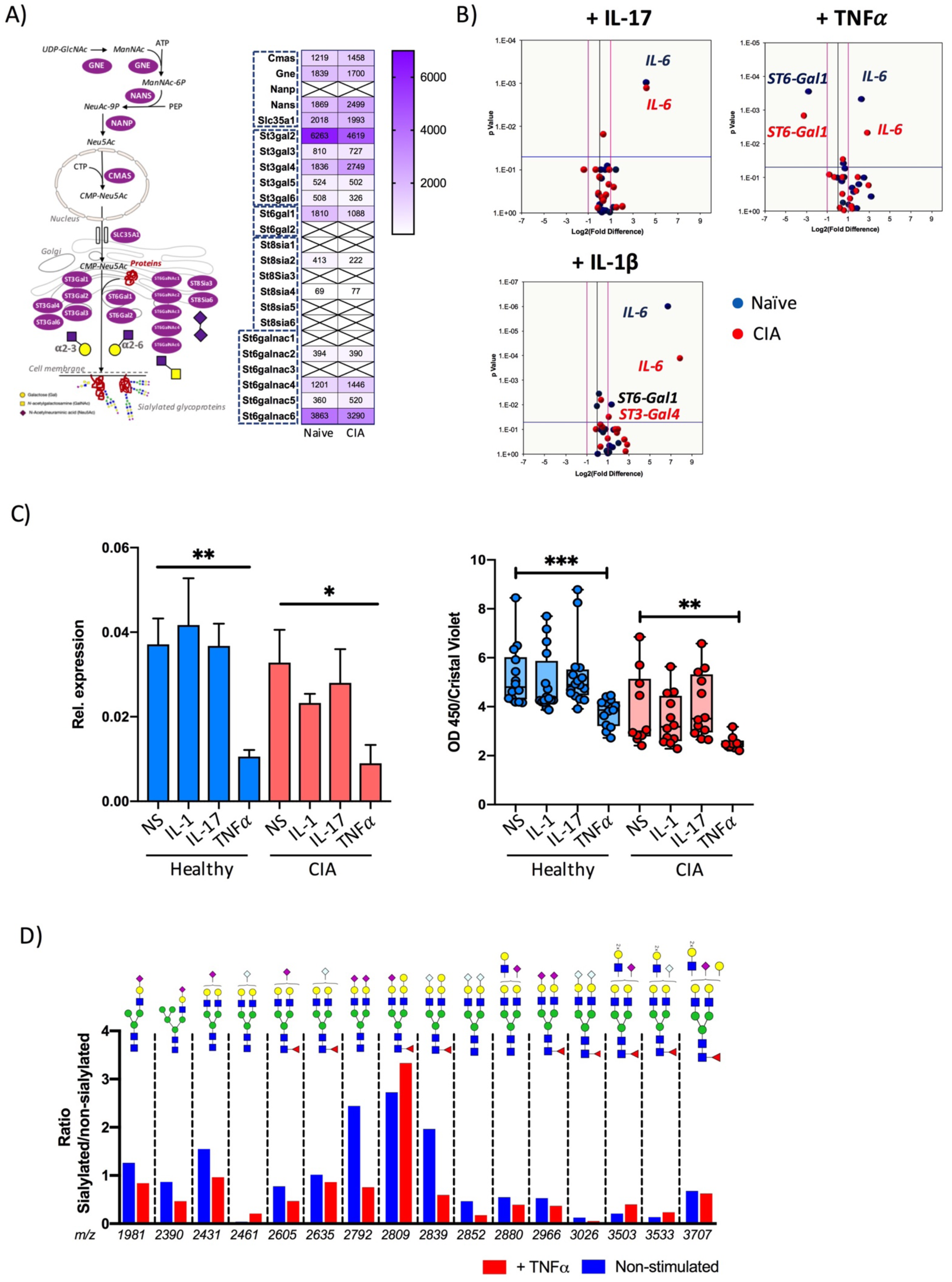
Down-regulation of α2-6 sialylation in arthritic synovial fibroblasts is mediated by TNFα-dependent inhibition of the sialyltransferase ST6Gal1. **a)** Synovial fibroblasts expanded from healthy or arthritic mouse joints were stimulated in vitro for 6 hours with recombinant IL-1β, TNFα or IL-17A [10 ng/ml]. Expression levels of genes involved in sialylation (as shown in diagram) in response to cytokine stimulation were evaluated by RT-qPCR. *IL-6* was included as a positive control to confirm cell activation. Volcano plots show log2 fold difference in stimulated cells (x axis) and p value (y axis). Each dot represents the mean of 3 independent experiments analyzed in triplicate, healthy cells in blue and arthritic cells in red. The pink lines indicate 2-fold-change in gene expression threshold and the blue line indicates the 0.05 threshold for the p value of the t-test used to evaluate statistical significance. **b)** TNFα, but not IL-1β or IL-17A reduces α2-6 sialylation of synovial fibroblasts both at RNA (*ST6Gal1* expression) and protein (SNA binding) levels. Expression of *ST6Gal1* (left panel) was evaluated by RT-qPCR as in (a). Results show the mean of 3 independent experiments analyzed in triplicate, error bars represent SEM. Statistical significance was determined using two tail unpaired t tests where *p < 0.05 and **p < 0.01. Expression of α2-6 sialylated glycoconjugates was determined by SNA binding as in Fig. 4a. Results show the merged results of 3 independent experiments, n=10-14. Statistical significance was determined using ordinary one-way ANOVA test, where **p < 0.01 and ***p < 0.001. c) Relative expression of individual sialylated glycan structures was evaluated by MALDI-TOF MS analysis as in Fig. 3. Ratios of sialylated vs non-sialylated twin structures were evaluated for non-stimulated cells (blue) and TNFα stimulated (red) cells (48 hours, 10ng/ml).

### Fibroblast populations are differentially sialylated in homeostatic and inflammatory conditions

The discovery of distinct subsets of SFs in different microdomains of the synovium has revolutionized our understanding of fibroblast biology in RA. Single cell transcriptomic experiments have shown that fibroblast subsets localize to specific regions in the synovium contributing to different aspects of disease pathogenesis^9,10^. For example, CD90+FAPα+ fibroblasts located in the synovial sub-lining are essential for perpetuation of the inflammatory response, whereas CD90-FAPα+ in the lining membrane are responsible for bone and cartilage damage^9^. Understanding which functional SF subset(s) lose sialic acid during inflammation would allow us to infer the dominant functional relevance of this glycan modification in SF-mediated pathogenesis. Therefore, we evaluated expression of ST6Gal1 and α2-6-Sialylation in CD90- and CD90+ SFs, representative of lining and sublining SFs respectively. As described previously, SNA and MAAII lectins were used to detect α2-6 and α2-3 sialic acid in SFs directly isolated from digested mouse synovium. PNA was used as a control, because sialic acid prevents its binding. SFs from healthy joints showed higher affinity for SNA than MAAII, supporting the homeostatic role of α2-6 sialylation. Interestingly, naïve, unstimulated CD90+ SFs had a higher basal α2-6 sialylation than CD90-SFs (Fig. 6a), perhaps suggesting that sialic acid content is related to functional SF specialization dependent on their anatomical location. We therefore FACS sorted CD90+ and CD90-SFs from naïve and CIA joints and levels of ST6Gal1, IL-6 and MMP3 mRNA were evaluated in both subsets (Fig. 6b). As expected, and further corroborating our results, ST6Gal1 was down-regulated in CIA SFs, but interestingly, ST6Gal1 was differentially down-regulated only in CD90+ SFs (Fig. 6b). MMP3 and IL-6 were also differentially expressed in both subsets (Fig. 6b), up-regulated during arthritis and preferentially expressed by CD90-SFs, consistent with findings from Croft et al ^9^and further demonstrating that CD90 discriminates functional subsets of SFs.

**Figure 6:**
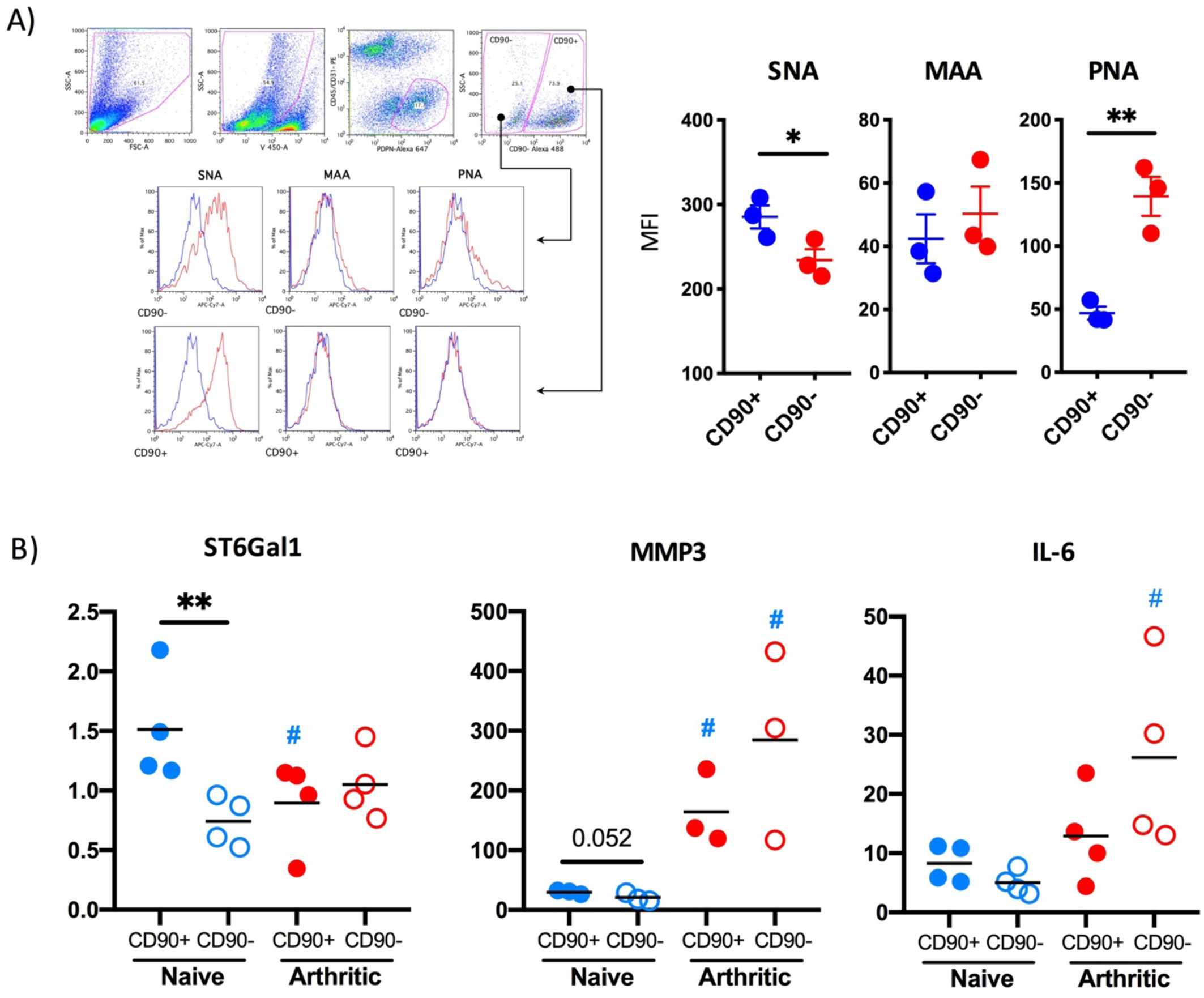
Reduction of α2-6 sialylation and ST6Gal1 levels is observed only in sublining CD90+ synovial fibroblasts. **a)** Synovial fibroblasts from healthy mice were isolated from mouse joints and identified by flow cytometry as in Fig. 1 (Zombie Violet-, CD45-, CD31-, Podoplanin+), separated into CD90+ and CD90-populations and stained with the biotinylated lectins SNA, MAAII and PNA and streptevadin-PE-Cy7. Graphs show the Mean Fluorescence Intensity for each cytokine and each dot represent cells from one individual mouse (n=3). b) Synovial fibroblasts from healthy and arthritic mice (n=4) were isolated from mouse joints as in (a) and RNA was purified using the RNeasy Mini Kit (Qiagen) according to manufacturer’s instructions. Expression of IL-6, MMP3 and ST6Gal1 mRNA was quantified by RT-qPCR. Statistics: one tail unpaired t-test was used for statistics, **p < 0.01, # p < 0.05 compared to the same population in control healthy mice.

### ST6Gal1 mediated sialylation of the SFs glycome is associated with disease remission

CD90+ SFs are differentially found in the sublining layer synovium, and are expanded in RA synovia immune responses^9,10,30^. As we observed that CD90+ SFs downregulate ST6Gal1 in inflammatory conditions, we postulated that a low sialylated SF-glycome would be a characteristic signature of environments undergoing strong immune reactions, as a consequence of high TNFα concentration in the synovial space. To test this hypothesis, we compared the N-glycome of SFs expanded from CIA animals, separating them into cells from non-affected paws (low scores, LS-SFs) and very inflamed paws (high scores, HS-SFs), taking advantage of the CIA model being asymmetrical, where not all limbs develop the same degree of inflammation. In addition, we included SFs from naïve non-arthritic mice. Unsupervised clustering of N-glycan relative expression, assessed by MS as before, revealed that one group of N-glycans that was strongly overexpressed in LS-SFs (*cluster 3619*, Fig. 7a). Strikingly, sialic acid was present in 11 out of 13 structures found within this cluster, consistent with idea that highly sialylated SFs is linked to a non-inflammatory phenotype. By contrast, *cluster 2080* including almost exclusively non-sialylated glycans was down-regulated (Fig. 7a). Because α2-6-linked sialic acid is a blocker for galectin binding and function ^31,32^, these findings suggested that a hypo-sialylated glycome, present in CD90+ SFs would be more sensitive to interactions with galectins by virtue of a higher exposure of galactosides, their natural ligand, being. Thus, we decided to stimulate naïve, LS- and HS-SFs (as in Fig. 7a) with recombinant galectin-3, described as a pro-inflammatory factor in RA ^33^, and IL-17 and TNFα as inflammatory mediators that do not bind glycans. Expression of IL-6 was used as a positive control to confirm SF activation, since it is a cytokine produced by SFs in response to the inflammatory cytokines used^19^. Indeed, all SF groups show some ability to respond to IL-17 and TNFα, being their response in direct correlation to their inflammatory status (Fig. 7b-c). However, LS-SFs did not respond to galectin-3, in clear contrast to HS-SFs (Fig. 7c) which were easily activated. This might be explained by the protective coating of sialic acid in the LS-SFs-glycome compared to HS-SFs, preventing galectin-3 binding and consequent inflammatory response. Nevertheless, LS-SFs still expressed elevated IL-6 expression compared with naïve SFs, and their glycome was significantly different to the naïve cells indicating that we cannot rule out that highly sialylated LS-SFs are in transition to become more inflammatory.

**Figure 7:**
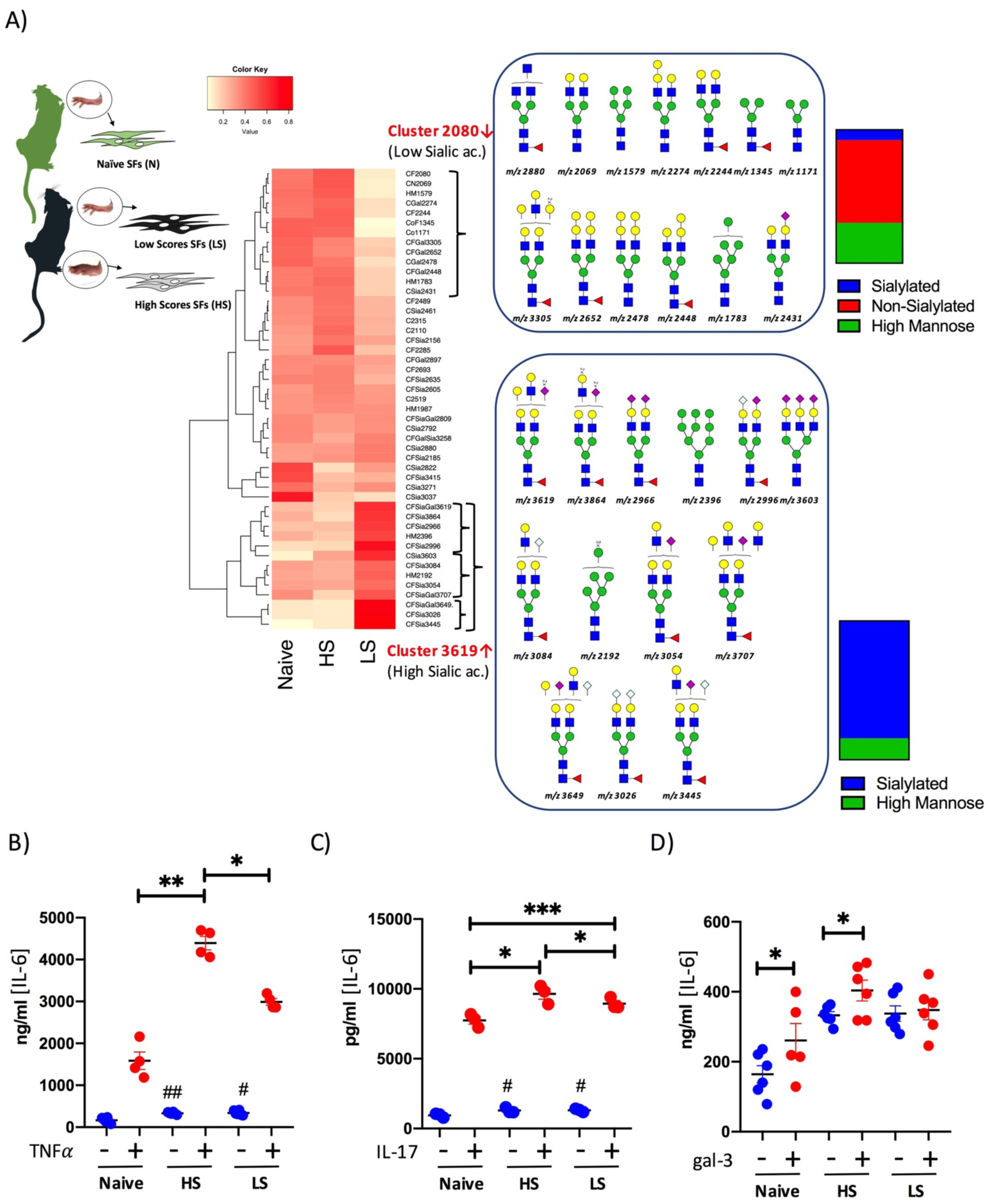
Synovial fibroblasts from arthritic mice show lower sialylation in non-inflamed joints than in highly inflamed joints. Synovial fibroblasts were isolated from healthy mice (naïve) and from CIA arthritic mice, where joints were separated into non-inflamed (Low scores, LS) and very inflamed (High scores, HS) prior to cell isolation and expansion. **a)** N-glycans were isolated and permethylated for to MALDI-TOF MS analysis and relative glycan expression was quantified as in Fig. 3. Two clusters were identified as distinctive of severe inflammation, for which glycan structures are shown. Percentage of sialylated and non-sialylated glycans are represented. Same cells from (a) were stimulated with TNFα **(a)**, IL-17 **(b)** [both at 10 ng/ml, 12 hours] or galectin-3 [1 ng/ml, 12 hours], when supernatants were collected and IL-6 levels were quantified by ELISA. Each dot represents an independent experiment analysed in triplicate, statistical significance was evaluated by ordinary one-way ANOVA and Tukey’s test for multiple comparations.

Hence, to assess whether high content of sialic acid is a protective marker, we used SFs expanded from either untreated early RA patients (<12 months from joint symptoms beginning), or RA patients in sustained remission under c-DMARDs+TNF-inhibitor combination therapy (Fig. 8). These samples were obtained from US-guided minimally invasive synovial tissue biopsies, which limited the amount of biological material and conditioned the coverage of MS-based studies. Nevertheless, we could still detect the most abundant structures, including high mannose and core fucosylated and non-fucosylated biantennary N-glycans. The ratio of mono and bi-sialylated versus their corresponding non-sialylated core provided us with the degree of sialylation in both clinical groups. Results showed that RA patients in sustained remission had a higher ratio of sialic-acid containing N-glycans, differences that were not seen when other non-sialylated structures were compared, such as ratios of fucosylated versus non-fucosylated or different mannose-containing glycans, suggesting that high sialic acid content is a distinctive feature of SF-glycome in RA patients in sustained remission (Fig. 8a). In line with this, SNA binding revealed a higher expression of α2-6 sialic acid in the synovium of RA patients in remission after c-DMARDs + TNF-inhibitor treatment (Fig. 8b) and in less inflammatory OA, supporting our conclusion of sialylation being an anti-inflammatory factor not only in the mouse model, but also in human RA.

**Figure 8:**
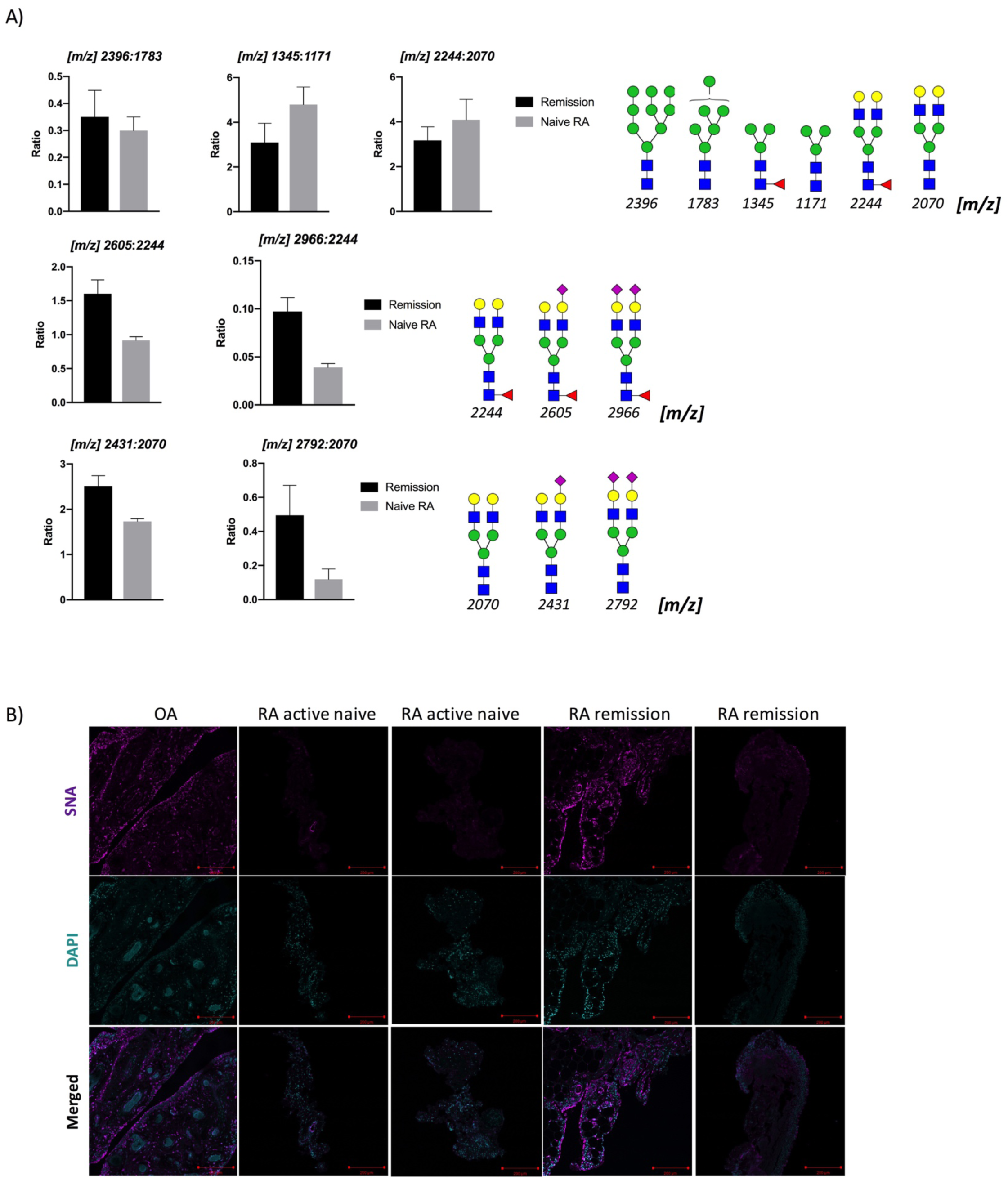
Levels of sialylation correlate with disease stages and pathotypes of human RA. **a)** Synovial fibroblasts were expanded from human biopsies from naïve RA patients (not exposed to any Disease Modifying Anti-Rheumatic Drug, n=2) and RA patients in sustained clinical and imaging remission achieved with c-DMARDs+TNFα-inhibitor combination therapy (n=3). N-glycans were isolated and subjected to MALDI-TOF MS analysis. Relative intensity was calculated, ratios of expression of the indicated paired glycans is shown. Glycan structures are designated by their m/z value. **b)** Synovium from naïve and remission RA patients was stained with biotinylated SNA and Alexa-647-conjugated streptavidin (magenta). DAPI (Cyan) was used to stain cell nuclei. Images were acquired with a confocal microscope Zeiss LSM 880. Each image is from one individual patient. Scale bars: 200 μm.

## Discussion

Glycans are involved in fundamental biological processes associated with SF-mediated pathophysiology, such as cell adhesion and migration, cell signaling and communication or immune modulation. One well described example of proteins recognizing glycans to initiate SF-dependent inflammatory responses is galectin-3, a carbohydrate binding protein that binds to galactoside-containing carbohydrates. Galectin-3 is a positive regulator of TLR-induced responses in human SFs^34^, it activates dermal and synovial fibroblasts to secrete pro-inflammatory IL-6, CXCL8 and MMP3, but it promotes a significantly higher secretion of TNFα, CCL2, CCL3, and CCL5 in SFs^19^, reflecting distinct stromal glycosylation and immunity of the synovial space. However, little attention has been given to the SFs glycome, the code read by carbohydrate binding proteins, such as galectin-3. Yang et al in 2004 aimed to characterize changes of surface glycoconjugates in bovine SFs upon TNFα and TGFβ stimulation^35^, being one of the first studies to evaluate SF glycosylation showing a link between cytokine(s) activity and changes in SFs-glycosylation. However, the structural information generated was very limited due to the nature of the lectin-based assays employed. More recent studies have focused on specific glycosyltransferases expressed in SFs, like fucosyltransferase 1 (Fut1) and galactosyltransferase-I (β1,4-GalT-I). Fut1 is involved in terminal α1-2 fucosylation and is up-regulated in RA synovial tissue^36^ regulating leukocyte-SF adhesion, whilst β1,4-GalT-I is induced by TNFα^37^ to promote binding to glycosylated extracellular matrix. Furthermore, aberrant glycosylation of fibronectin in RA has also been reported^38^. Nonetheless, the nature of the SF-glycome as a whole is unknown, and the role of glycosylation in SFs during joint inflammation is not understood. Therefore, we aimed to conduct a broader systematic study to bridge the gap between glycomics and other omics approaches in the field of SF-dependent immunology. Our results here provide an extensive description of the SF glycome in heath and disease that will help to further understand the interactions of SFs with other immune cells and matrix components in the joint tissue.

The success of glycosciences in the field of cancer is an exemplar model that integrates glycomics into stromal immunology. Distinct modifications in tumors include truncated N- and O-glycans, increased N-glycan branching and changes in sialylation. Ultimately, these changes affect fundamental processes of cancer biology like cell adhesion, signaling, tumor progression and metastasis^23^, making glycans attractive targets for therapeutic intervention. Recent technological advances and the prospective use of glycan-based biomarkers for patient stratification have led it to be estimated that the global glycobiology market size to grow 14.7% in the next years, with an estimated market value of USD 822.5 million in 2018. In RA, SFs undergo important epigenetic changes to adopt a tumor-like invasive phenotype that are also heavily determined by glycosylation including hyperproliferation, secretion of MMPs responsible of tissue damage and migration, and perpetuation of local proinflammatory responses. Therefore, it is likely that a better understanding of SF-glycobiology aids the identification of novel glycan-based therapeutics to target SFs, mirroring recent advances in cancer and opening new therapeutic opportunities for patients that are refractory to immunosuppressive drugs.

We combined transcriptomic and MS-based glycomic analysis to define changes in SF-glycome with unprecedented detail in synovial inflammation. Our results show that N-glycome from healthy SFs comprises LacNAc-containing structures with high levels of core fucosylation and terminal sialylation. However, arthritis renders the sublining CD90+ SF population hyposialylated, specifically in α2,6-linkage, which would increase the accessibility of LacNAc repeats, allowing galectin-3 binding to SF surface glycoconjugates. In fact, sialylation can act as a ‘switch off’ for galectin-3 function. For example, ST6Gal1 up-regulates α 2-6 sialylation to block binding of galectin-3 to β1 integrin, inhibiting cell apoptosis^39^. Sialylation also inhibits galectin-3 binding to squamous epithelia and tumour cells ^40-42^. Thus, because of a desialylated glycome, galectin-3 could induce secretion of inflammatory mediators such as IL-6 or Ccl2 in SFs, increasing local concentrations of TNFα that would further reduce ST6Gal-1 and α2-6 sialylation, establishing an autocrine pro-inflammatory loop that could explain perpetuation of disease in a similar fashion shown by members of the IL6 family^43^. By contrast, healthy synovium shows higher levels of α2-6-Sialylation in SFs, perhaps masking LacNAc repeats and preventing inflammatory actions of galectins. Thus, we propose that sialylation might be a homeostatic mechanism that is lost during RA progression because of TNFα overexposure.

To understand the physiological role of sialic acid in SFs, the role of carbohydrate binding proteins (CBPs) requires consideration. CBPs are expressed either in SFs, or in other immune cells, promoting trans or cis interactions. Siglecs (sialic acid-binding immunoglobulin-type lectins) are a family of immune regulatory proteins primarily found on hematopoietic cells. Consistent with our proposed homeostatic role for sialic acid in the synovium, Siglec-9 protected mice against experimental arthritis, although authors reported that it had no effect on SFs^44^. Instead, Siglec-9 inhibited NF-kB activation in human RA macrophages. However, effects on SFs cannot be ruled out, as following cytokine stimulation during disease, RASFs might already have a reduced sialylation profile which would make them unresponsive to Siglec-mediated actions. It might therefore be possible that siglecs mediate regulatory effects in synovial self-tolerance under healthy conditions, or at very early disease stages. The mechanism could involve Siglec-sialic acid trans interactions between SF and immune cells. In line with this, B cells lacking Siglec-2 (CD22) and Siglec-G develop spontaneous autoimmunity^45^ and Siglec-G^-/-^ lupus-prone MRL/lpr mice exhibit increased severity and early onset of arthritis^46^. Besides, most siglecs (2,3,5,6-11) show inhibitory effects on TLR-dependent activation and mediate immunosuppression in the tumor microenvironment because of local hypersialylation^47^. By contrast, Siglec-1 (sialoadhesin) shows pro-inflammatory actions, it is up-regulated in activated macrophages in RA^48^ and suppresses Tregs^49^. Interestingly, the anti-inflammatory Siglec-2 has predilection for α2-6 sialic acid, but the pro-inflammatory Siglec-1 binds preferentially to α2-3^50^, thereby supporting our conclusion that pro-inflammatory SFs diminish α2-6 sialylation but not α2-3. Thus, homeostatic α2-6>>>α2-3 sialylation on SFs could induce Siglec-2-mediated signals to prevent B cell activation, whereas inflammatory α2-6<α2-3 sialylation would support SFs interactions with pathogenic macrophages via Siglec-1, cells that also secrete high levels of pro-inflammatory galectin-3 and TNFα, thereby perpetuating disease. Although this is an oversimplification of a complex scenario, it provides a good example of how the axis cytokines-sialylation-siglecs/galectins could regulate interactions between immune cells and SFs. Moreover, other sialic acid-binding proteins could be relevant, like selectins. Selectins are expressed in lymphoid, myeloid and endothelial cells, and mice deficient in Selectin-P and selectin-E show an enhanced arthritis progression^51,52^. Conversely, ficolins, innate immune receptors recognizing sialic acid too, show opposite effects, as ficolin B deficient mice are protected against arthritis^53,54^. These glycan-dependent networks and associated signaling for each sialic acid binding protein could explain the SF-mediated actions to establish chronic inflammation and interactions with immune cells in the arthritic synovium. Thus, further work is necessary to determine expression of these sialic acid-binding proteins in different cell types in the synovium to complement the findings presented in this work.

These cellular networks will be further defined by inclusion of additional immune cells in the equation, but also by the SFs anatomical and functional heterogeneity that has been recently revealed^10,30^. Distinct CD90+ SFs subsets found in the sublining synovial areas are expanded in RA synovium and are indeed the main contributors to inflammatory responses, results that have been confirmed in animal models ^9^. Our results in the mouse model indicate that only sublining CD90+ SFs reduce ST6-Gal1 expression, compared to the lining CD90-SF suggesting that sialic acid would exert different effects on the SFs subsets interacting with other immune cells, which could offer potential pathways to modulate specific immune networks in the synovium. We can also hypothesize that these glycan-dependent interactions could explain the observed disease tissue heterogeneity described in RA patients, including lymphoid, myeloid and fibrotic phenotypes^55,56^. Because synovial phenotypes observed in RA have been associated to distinct pathways (myeloid-IL-1β/TNF, lymphoid-IL-17^55^, it would be relevant to study ST6Gal1 expression to assess whether pro-inflammatory sialylation correlate with synovium RA phenotypes. Interestingly, TNFα induces a similar shift of α-2,6/α-2,3-sialylation in chondrocytes downregulating ST6Gal1^57^, in agreement with the pro-inflammatory role of α2,3-sialylation in synovial joints.

In conclusion, in the present report, we revealed that the reduction of α2,6-linked terminal sialylation constitutes a signature of the inflammatory, vs damage status of synovial fibroblasts. This altered phenotype seems to be induced by TNFα, in contrast with IL-1 or IL-17 that had no effect. This might open a possibility to segregate disease phenotypes by virtue of their glycosylation status. Future studies clarifying the functional consequences of shifting sialylation patterns in SFs are now required, as well as expression of sialic acid receptors in other cells, like B cells or monocytes. The vast, and yet unexploited amount of information contained in the SF glycome could offer novel therapeutic targets, and further work is anticipated before this goal can be achieved. Nevertheless, this study strengths the foundations for future clinic intervention of glycan-dependent pathological networks in RA, and other autoimmune diseases where stromal fibroblasts control local inflammation.

## Figure legends

**Supplemental Figure 1:**
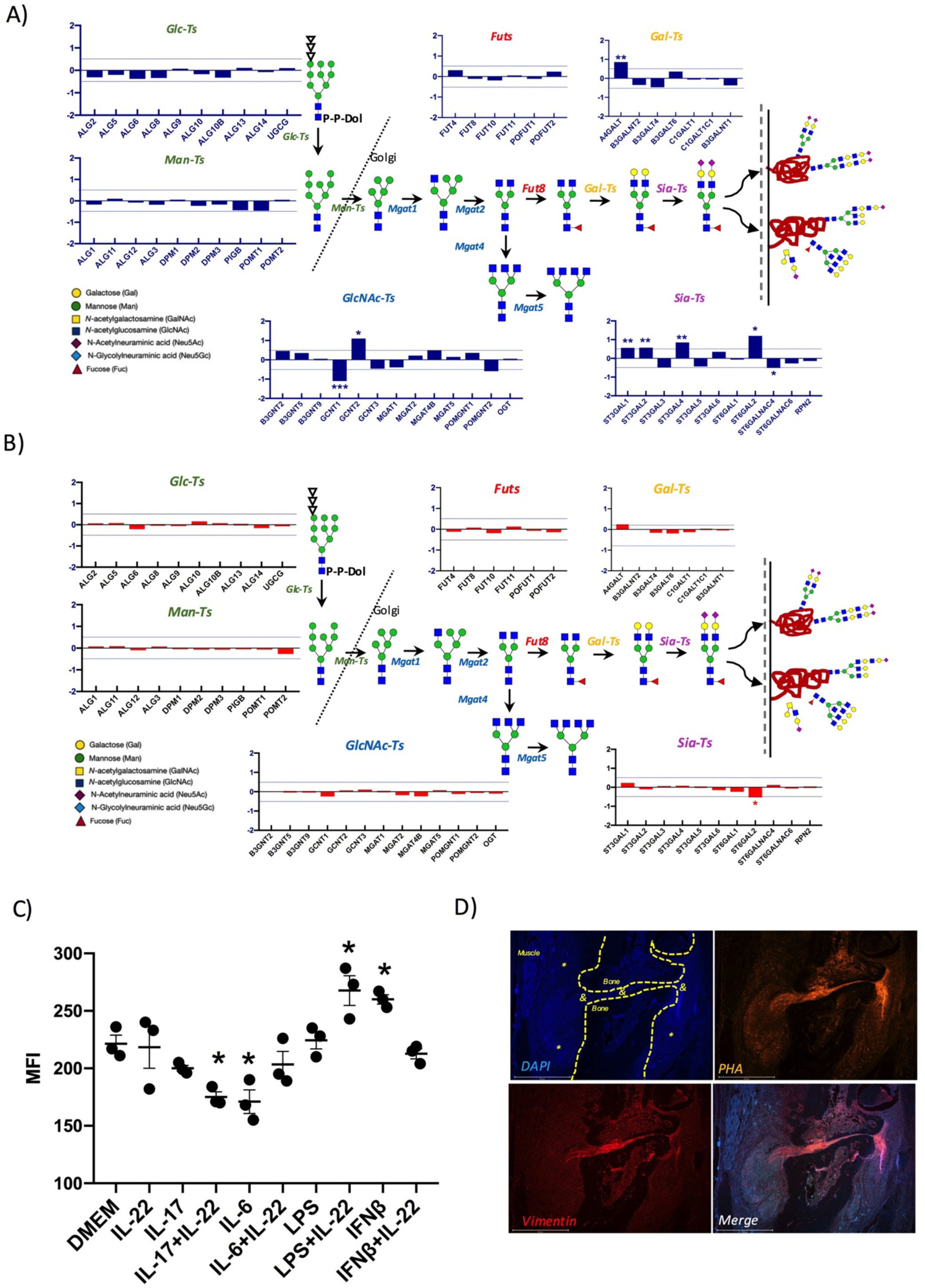
Cytokines regulate glycosylation in synovial fibroblasts. **a-b)** Gene expression of human SFs was analyzed following stimulation with TNFα, 1 ng/ml, 6h (a) and IL-17, 10 ng/ml, 6h (b), data from Slowikowski et al ^25^. Expression of glycosyltransferase genes was analyzed in https://immunogenomics.io/fibrotime, *p < 0.05 and **p < 0.01 **c)** Murine SFs were isolated and expanded from mouse synovium and incubated with the indicated inflammatory mediators (10ng/ml for IL-22, IL-17, IL-6, and IFNβ and 0.2 μg/ml for LPS) for 24 hours, when binding of PHA was evaluated by Flow Cytometry. Statistical significance was determined using ordinary one-way ANOVA with Bonferroni’s multiple comparisons test, *p < 0.05. **d)** Joints of CIA mice show an association of PHA binding (multi-branched glycans) and stromal cells in inflamed areas. Bone is shown with dotted yellow lines, (*) immune cell infiltration, (&) stromal-mediated inflammation. Stromal marker vimentin (red), PHA binding (orange) and DAPI staining (blue) are shown.

**Supplemental Figure 2:**
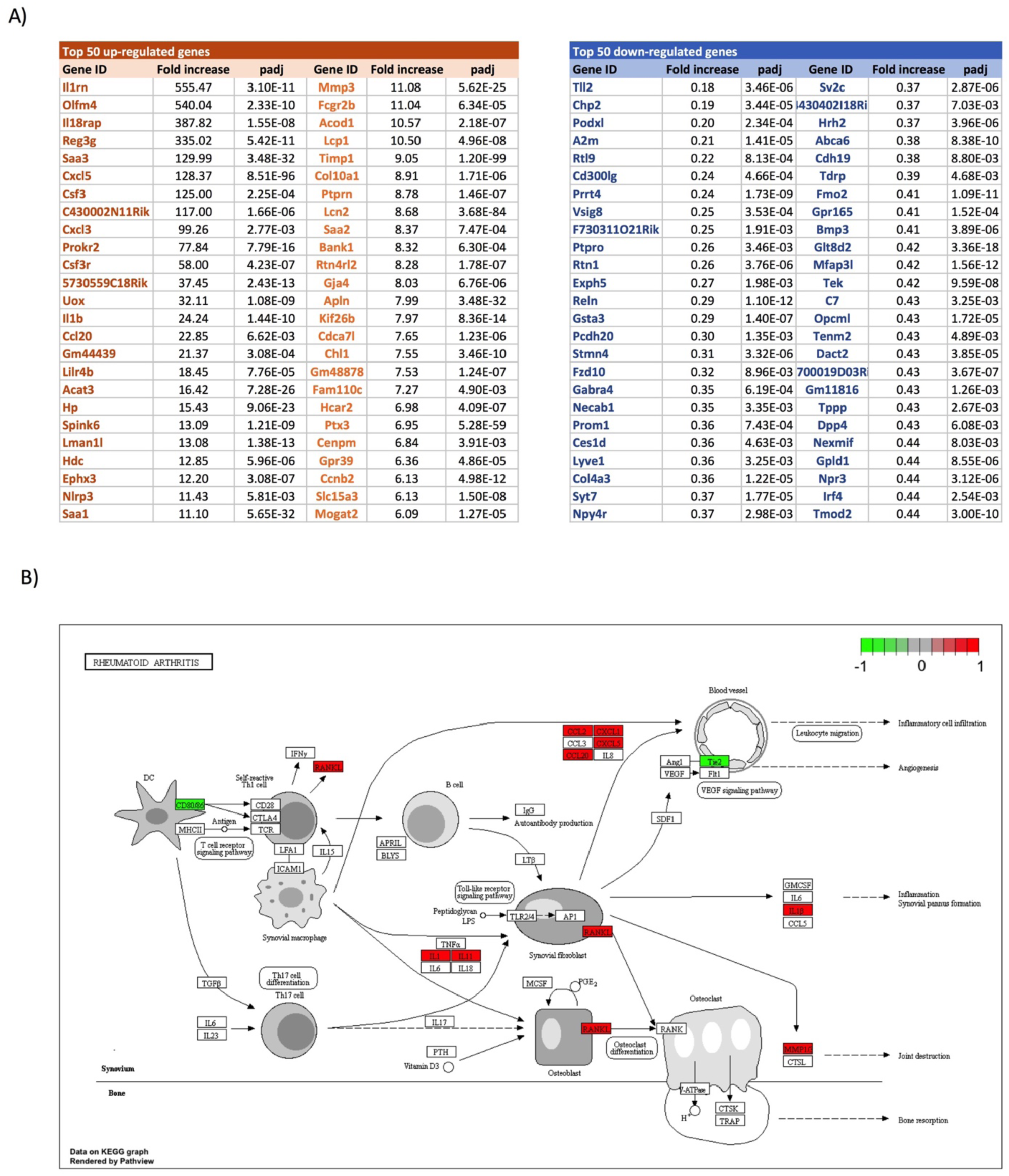
Rheumatoid Arthritis disease pathway as defined by KEGG database. DE genes in CIA SFs compared to healthy controls are shown in red (up-regulated) or green (down-regulated).

**Supplemental Figure 3:**
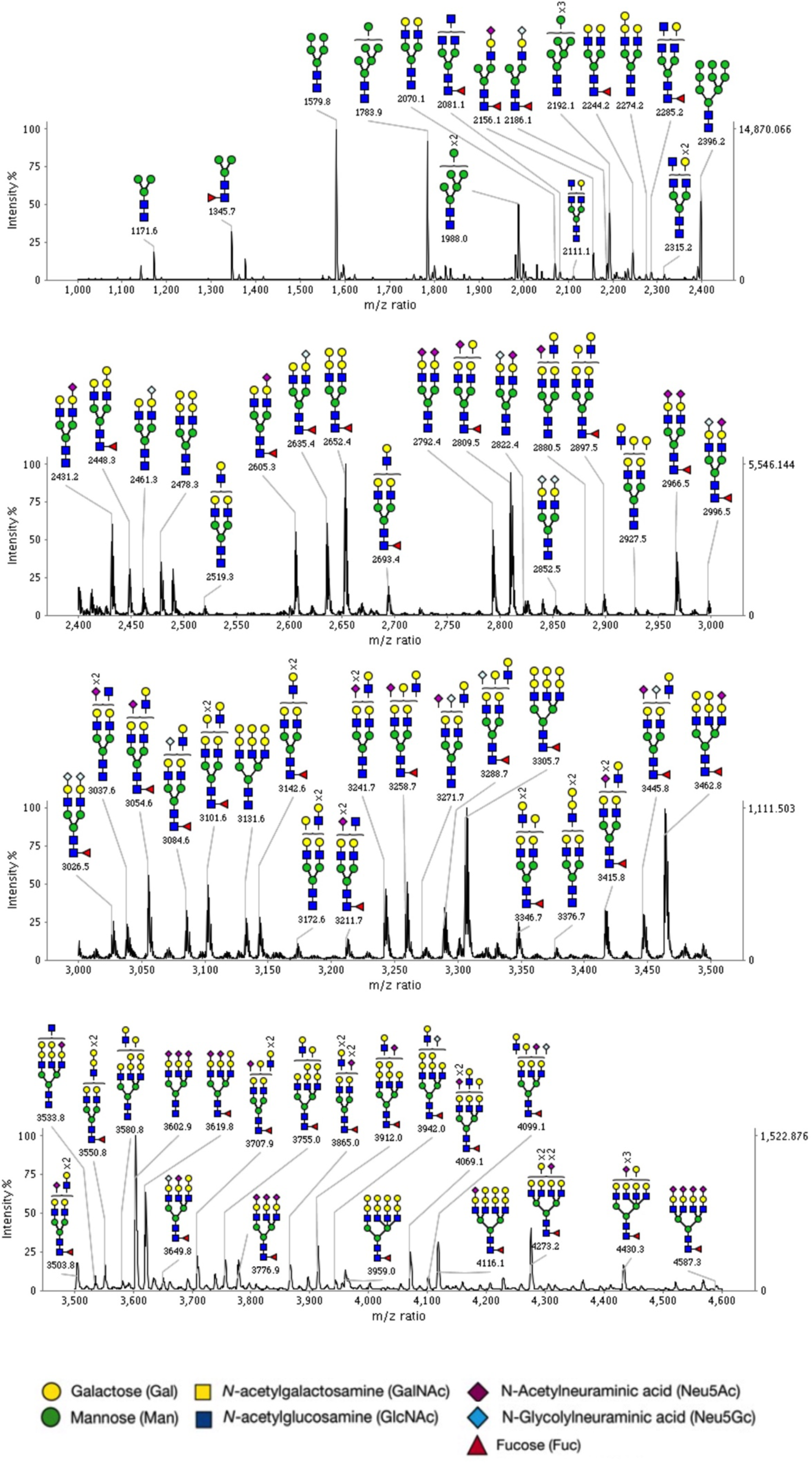
MALDI-TOF MS profiles of the permethylated N-linked glycans derived from expanded murine SFs. Data were obtained from the 50% MeCN fraction, and all molecular ions are present in sodiated form ([M + Na]+).

**Supplemental Figure 4:**
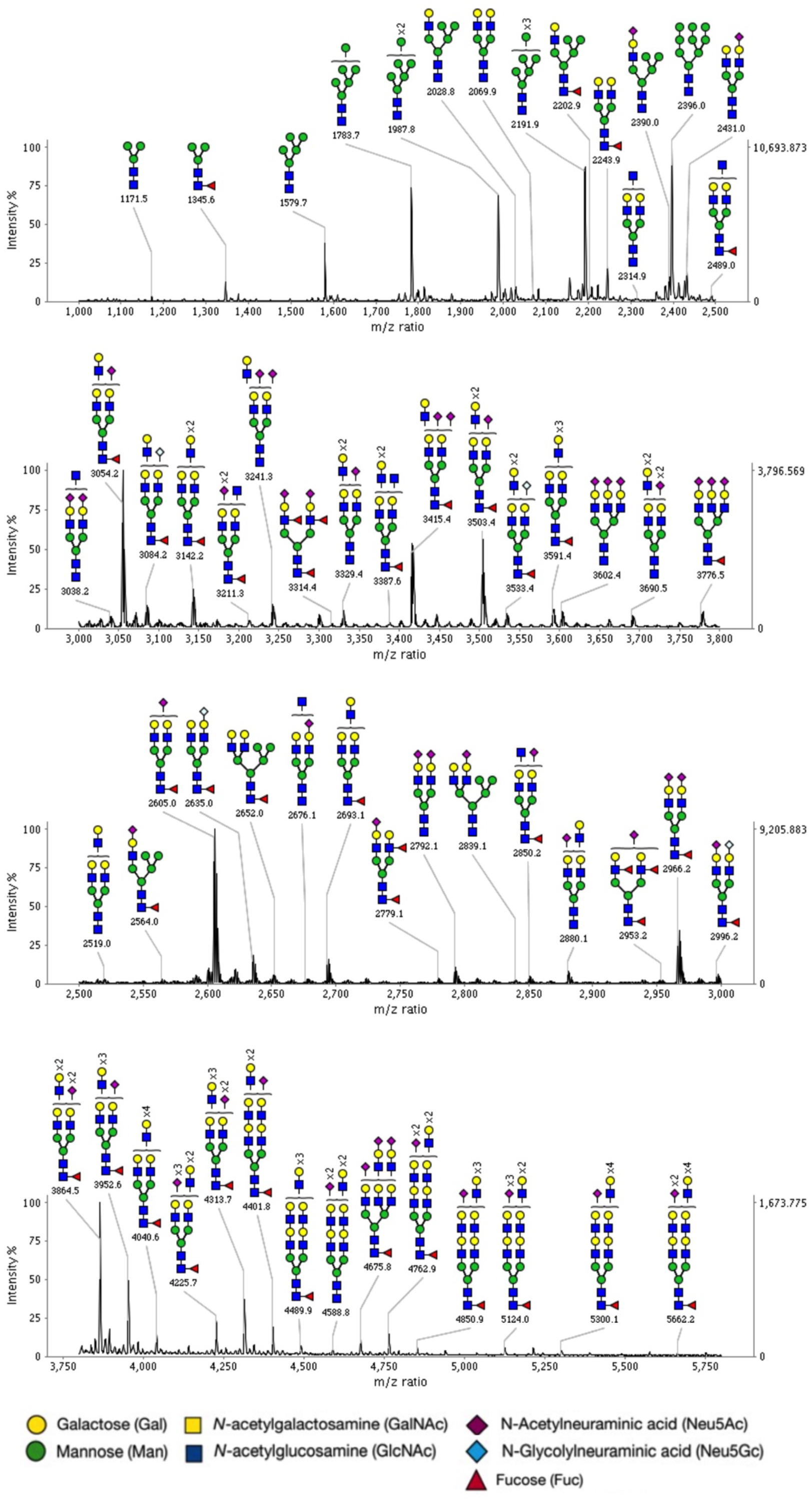
MALDI-TOF MS profiles of the permethylated N-linked glycans derived from expanded human SFs. Data were obtained from the 50% MeCN fraction, and all molecular ions are present in sodiated form ([M + Na]+).

## Methods

### Patient recruitment and isolation of human fibroblasts lines

Tissue samples were collected from patients who fulfilled the American College of Rheumatology (formerly, the American Rheumatism Association) 1987 revised criteria for RA ^58^. Tissue samples were also acquired from patients with radiographically confirmed OA. Synovium samples and overlying skin were obtained from the knee joint of each patient at the time of joint replacement surgery. This study was reviewed and approved by the South Birmingham Local Ethics Committee (LREC 5735), all patients gave written informed consent. Tissue samples were minced into 1 mm^3^ sections under sterile conditions, washed, and then resuspended in 10 ml 5 mM EDTA in phosphate buffered saline (PBS) and incubated for 2 hours at 4°C with vigorous shaking. The resulting cell–tissue mixture was washed 3 times in RPMI medium and then cultured in complete RPMI 10% FCS until adherent fibroblast colonies became confluent. Non-adherent cells and tissue fragments were discarded once adherent cells had appeared. Fibroblasts were expanded until reaching confluence and then reseeded into tissue culture flasks of twice the surface area after trypsin treatment. All experiments using expanded fibroblast lines used cells between passages 3 and 4. For active naïve and remission RA patients^59^, biopsies were collected using a standard operating procedure with sampling of at least 6-8 different tissue pieces as required by the international recommendations for synovial tissue analysis^60,61^.

### Immunofluorescence analysis

10,000 cells were grown in chamber slides, wash 3 times with PBS and fixed with 4% PFA for 20 minutes at room temperature. Cells were permeabilized with PBS 0.5% Triton and rinsed with PBS-Tween 20. Endogenous biotin was blocked with the Streptavidin/Biotin blocking kit according to manufacturer’s instructions (Vector Laboratories, SP-2002). Carbo-Free Blocking Solution (Vector laboratories, SP-5040) was used as blocking solution. Samples were incubated overnight with biotinylated lectins at 2 µg/ml in PBS purchased from Vector laboratories [Concanavalin A (ConA), B-1005; Ricinus Communis Agglutinin I (RCA), B-1085; Peanut Agglutinin (PNA), B-1075; Jacaline, B-1155; Wheat Germ Agglutinin (WGA), B-1025; Sambucus Nigra Lectin (SNA), B-1305; Lycopersicon Esculentum Lectin (LEL), B-1175; Erythrina Cristagalli Lectin (ECA), B-1145; Aleuria Aurantia Lectin (AAL), B-1395; Ulex Europaeus Agglutinin I (UEA), B-1065]. Lectins were finally detected with FITC-conjugated streptavidin (Biolegend, 405201). Mouse joint sections (7 μm) and human synovium sections were deparaffinized in xylene and dehydrated in ethanol, and antigen was retrieved by incubation at 60°C for 40 minutes in sodium citrate buffer (10mM Sodium Citrate, 0.05% Tween 20, pH 6.0). Blocking was performed as explained before. Samples were staining with biotinylated Phaseolus Vulgaris Leucoagglutinin (PHA-L) from Vector laboratories, catalog number B-1115 and Alexa-555-cojugated streptavidin (Thermofisher, S21381). Vimentin was detected with anti-vimentin goat IgG (Sigma, V4613) and Alexa-647 conjugated anti goat IgG (Thermofisher, A21245). All samples were counterstained with DAPI. Images were obtained using an LSM 510 META confocal laser coupled to an Axiovert 200 microscope (Zeiss) and analyzed with Zeiss LSM Image Browser software.

### Flow cytometry

Cultured cells were detached from the plates using Accutase cell detachment solution (Stemcell Technologies) according to manufacturer’s instructions. For lectin staining (listed before), cells were blocked with Carbo-Free Blocking Solution (Vector laboratories, SP-5040) and stained with biotinylated lectins and FITC-conjugated streptavidin in PBS for 20 minutes at 4C. Data were acquired using a BD LSRII flow cytometer and analysed with FlowJo software.

### Mice and Collagen-Induced Arthritis (CIA) model

CIA was induced in 8–10-week-old male DBA/1 mice (Envigo) on day 0 by intradermal immunization with bovine type II collagen (MD Biosciences) in Freund’s complete adjuvant (CFA). At day 21, mice received 100 mg of collagen in PBS intraperitoneally. Disease scores were measured every 24 hours in a scale from 0-4 for each paw. Animals reaching an overall score of 10 or more were immediately euthanized. Animals were maintained in the University of Glasgow Biological Services Units in accordance with the Home Office UK Licenses P8C60C865, I675F0C46 and ID5D5F18C and the Ethics Review Board of the University of Glasgow.

### Isolation of murine synovial fibroblasts and explant cell culture

For mouse synovial tissue digestion, skin and soft tissues were removed from mouse limbs and bones with intact joints were dissected and transferred into DMEM (+5% FCS) containing 1% L-Glutamine, 1% Penicillin Streptavidin, nystatin and 1 mg/ml type IV collagenase (Sigma) and 5 mg/ml DNase I. Samples were incubated in the shaking oven at 37 °C for 1:20 h, when EDTA was added, final concentration 0.5 mM for 5 minutes at 37 °C. Samples were vortexed to release cells. Cells were centrifuged and washed with DMEM 10% FCS twice. For cell expansion, cells were plated and expanded until adherent fibroblast colonies became confluent. Non-adherent cells and tissue fragments were discarded once adherent cells had appeared, usually after 48 hours. Fibroblasts were expanded until reaching confluence and then reseeded into tissue culture flasks after trypsin treatment. Expanded synovial fibroblasts were used at passage 3 or 4 only, when culture purity was assessed by flow cytometry and expression of CD106 (Biolegend, 105722), CD54 (Biolegend, 116107), and CD90 (Biolegend, 105316). Expression of CD11b was also evaluated (Invitrogen, 11-0112-85), where cells were <1% positive.

### Mouse synovial fibroblasts sorting

Single cell suspensions from mouse synovium were obtained as described before. Cells were stained at 4 °C with Zombie Violet staining (BioLegend, 423113) to exclude dead cells. Antibodies used were anti-CD45 (Biolegend, 103106), anti-CD90 (Biolegend, 103106), anti-podoplanin (Biolegend, 105316), anti-CD31 (Invitrogen, 12-0311-81), Cell sorting was performed immediately after staining using a FACS Aria Ilu machine. For sorted populations, purity was determined by reanalysis for the target population based on cell surface markers immediately post sorting. Purity was >99% for the synovial fibroblasts target population (CD31-, CD45-, podoplanin+).

### RNA-Seq preparation, sequencing and data analysis

Total RNA from sorted synovial fibroblasts was isolated immediately post sorting using RNeasy Micro kit (Qiagen, Germany). RNA integrity was checked with the Agilent 2100 Bioanalyzer System. All purified RNA had a RIN value >9. Libraries were prepared using the TruSeq mRNA stranded library preparation method. Samples were sequenced 2×75bp to an average of more than 30 million reads. Kallisto was used to quantify the transcript abundance of RNA-seq reads. All RNAseq reads were then aligned to mouse reference genome (GRCM38) using Hisat2 version 2.1.0. Featurecounts version 1.4.6 was used to quantify reads counts. Data quality control, non-expressed gene filtering, median ratio normalization (MRN) implemented in DESeq2 package and identification of differentially expressed (DE) genes were done using the R bioconductor project DEbrowser ^62^. Genes that passed a threshold of padj < 0.01 and log2foldChange > 2 in DE analysis were considered for further analysis. Gene Ontology (GO) enrichment, KEGG pathway enrichment and UniProt Keywords enrichment were performed in String (https://string-db.org) based on statistically significantly DE genes.

### Mass spectrometric analysis of synovial fibroblasts N-glycans and unsupervised hierarchical clustering

Synovial fibroblasts cells were scraped off tissue culture plates and suspended in iced-cold ultrapure water before homogenization and sonication were performed. Cells protein extract was precipitated in a methanol/chloroform extraction, Cell extracts were reduced and carboxymethylated, using Dithiothreitol and Iodoacetic acid, and then treated with trypsin. The treated samples were purified using a C18 cartridge (Oasis HLB Plus Waters) prior to the release of N-glycans by PNGase F (recombinant from Escherichia coli, Roche) digestion. Released N-glycans were permethylated and then purified using a Sep-Pak C18 cartridge (Waters) prior to MS analysis. The resulting enzyme-treated samples were lyophilized and permethylated prior to MS analysis. Purified permethylated glycans were dissolved in 10 µl methanol and 1 µl of the sample was mixed with 1 µl of matrix, 20 mg/ml 2,5-dihydroxybenzoic acid (DHB) in 70% (v/v) aqueous methanol and loaded on to a metal target plate. 4800 MALDI-TOF/TOF mass spectrometer (AB SCIEX) was run in the reflectron positive ion mode to acquire data. MS spectra were annotated manually with the assistance of the glycobioinformatics tool GlycoWorkBench (GWB). All N-glycans were assumed to have a Manα1–6(Manα1–3)Manβ1– 4GlcNAcβ1–4GlcNAc core structure based on knowledge of the N-glycan biosynthetic pathway in mammalian cells and PNGase F specificity. The composition of the glycans derived from MALDI-TOF MS in positive ion mode were manually interpreted. Relative expression of individual glycan structures was evaluated by calculating areas under the curve of annotated peaks using GWB. For comparative studies, MS peak areas were quantified and normalized against total measured intensities. Unsupervised hierarchical clustering with Euclidean distance was performed on the relative glycan expression samples using the heatmap.2 function of the gplots package in R. The symbolic nomenclature used in the all the spectra annotations is the same as the one used by the Consortium for Functional Glycomics (CFG) (http://www.functionalglycomics.org/).

### Enzyme-linked lectin assays (ELLA) and ELISA

Synovial fibroblasts at 10.000 cells/well were grown on 96-well plates. Cells were washed three times with cold PBS. Cells were then fixed with 4% paraformaldehyde for 20 min. Fixed cells were washed three times with PBS containing 0.05% Tween. CarboFree solution (Vector Laboratories) was used to block non-specific interactions, following incubation with 1 µg/mL of biotinylated lectin (Vector Laboratories, Burlingame), for 30 min. Cells were washed with PBS and incubated with HRP conjugated streptavidin for 20 minutes, and washed again with PBS-Tween. A reaction was induced in the cells with developing solution, consisting of 1 mg/mL p-nitrophenyl phosphate in 0.5 mmol/L. The reaction was allowed to proceed in the dark and plates were read at 405 nm using a microplate spectrophotometer. Interleukin-6 (IL-6) expression was measured by ELISA in SF-supernatants according to the manufacturer’s instructions (BD Biosciences, Oxford, UK).

*qRT-PCR*. Cells were lysed in RLT lysis buffer prior to mRNA extraction using RNeasy Plus Mini kit (Qiagen, Germany) according to the manufacturer’s instructions. The High Capacity cDNA Reverse Transcriptase kit (Applied Bio-systems, Life Technology, UK) was used to generate cDNA for use with StepOne PlusTM real-time PCR system (Applied Biosystems, UK) and predesigned KiCqStart® qPCR Ready Mix (Sigma-Aldrich) or TaqMan gene expression assays (Applied Biosystems). Pre-designed KiCqStartTM primers (Sigma-Aldrich) were purchased to evaluate β-Actin, Cmas, Gne, Nanp, Nans, Slc35a1, ST3Gal2, ST3Gal3, ST3Gal4, ST3Gal5, ST3Gal6, ST8Sia1, ST8Sia2, ST8Sia3, ST8Sia4, ST8Sia5, ST8Sia6, ST6GalNAc2, ST6GalNAc4, ST6GalNAc2, ST6GalNAc4, ST6GalNAc5 and ST6GalNAc6. TaqMan predesigned probes (Thermofisher Scientific) were used to evaluate Actin, IL-6, ST6Gal1, ST6Gal2, ST3Gal1, ST6GalNAc1, ST6GalNAc1, ST6GalNAc3 and MMP3. Data were normalised to the reference gene β-actin to obtain the ΔCT values that were used to calculate the fold change from the ΔΔCT values following normalisation to biological control group.

## Acknowledgements

The work was funded by an award to M.A.P. from Versus Arthritis (21221).

